# Interferon Alpha and Kinase Inhibitor Nilotinib Increase Cell Adhesion and Tunneling Nanotubes in CML

**DOI:** 10.1101/297838

**Authors:** Maria Omsland, Vibeke Andresen, Pilar Ayuda-Durán, Stein-Erik Gullaksen, Randi Hovland, Jorrit Enserink, Bjørn Tore Gjertsen

**Affiliations:** Centre for Cancer Biomarkers (CCBIO), Department of Clinical Science, University of Bergen, Bergen, Norway; Department of Internal Medicine, Haukeland University Hospital, Bergen, Norway; Department of Molecular Cell Biology, Institute for Cancer Research, The Norwegian Radium Hospital, Oslo University Hospital, Montebello, Oslo, Norway; Department of Medical Genetics, Haukeland University Hospital, Bergen, Norway.

**Keywords:** (5) Chronic myeloid leukemia (CML), Tunneling nanotubes (TNTs), Tyrosine Kinase inhibitors (TKIs), Interleukins, Cell adhesion

## Abstract

**Summary statement:** This study describes the effects of tyrosine kinase inhibitors on tunneling nanotube formation via increased adhesion through β-integrin in chronic myeloid leukemia cells.

**Abstract:** The actin-containing cell-to-cell communicator tunneling nanotube (TNT) is involved in regulation of cell death threshold of leukemic cells, while the mechanism of TNT regulation is mostly unknown. We have investigated TNT formation and its response to treatment in chronic myeloid leukemia (CML) cells with the pathognomonic chimeric fusion kinase BCR-ABL1 after treatment with the tyrosine kinase inhibitor nilotinib and interferon-α. Bone marrow cells of chronic phase CML patients and the CML cell line Kcl-22 formed few or no TNTs. Nilotinib and interferon-α treatment induced TNT formation in Kcl-22 cells and were found to be linked to increased adherence to fibronectin coated surfaces by restoration of β1-integrin function. This suggests modulation of TNT cell-cell communication in CML as a novel mechanism in kinase inhibitor therapy of CML.

## Introduction

Chronic myeloid leukemia (CML) is a myeloid stem cell disease characterized by the BCR-ABL1 fusion protein derived from the chromosomal translocation t(9;22), involving bone marrow and spleen in the chronic phase. The role of BCR-ABL1 in impaired communication between cells in the microenvironment (Bhatia et al., 1995; Gordon et al., 1987) is less understood in the context of the efficient therapies with small molecule kinase inhibitors that emerged at the millennium (Bruck et al., 2018; Hochhaus et al., 2017).

The BCR-ABL1 protein has a filamentous (F)-actin binding domain and orchestrates several cellular processes involving actin processing, cell attachment to fibronectin and cell migration (Wertheim et al., 2003). Features of CML progenitor cells from patients in the chronic phase include increased motility and low affinity to fibronectin coated surfaces compared to normal counterparts (Verfaillie et al., 1992). Interferon alpha (IFNα), previously pivotal in CML therapy, increase adhesion of CML progenitor cells to bone marrow stromal cells (Dowding et al., 1991). Attenuated cellular mobility seems therefore to be a significant mechanism of action in effective CML therapy, recently revisited in the effective therapeutic combination of a tyrosine kinase inhibitor (TKI) and IFNα eradicating CML progenitor cells resulting in non-detectable disease (Hjorth-Hansen et al., 2016; Simonsson et al., 2011).

It is well established that the tumor microenvironment and cell-cell interaction plays a pivotal role in the outcome of cancer therapy (Joyce and Pollard, 2009). One such form of physical interaction is the tunneling nanotube (TNT) (Rustom et al., 2004). TNTs are defined as thin (50-200 nm), fragile and dynamic structures, consisting of plasma membrane and F-actin (Abounit and Zurzolo, 2012; Rustom et al., 2004). They are involved in cell-cell interaction and intercellular transport of organelles and pathogens such as virus and bacteria (Gousset et al., 2013; Gurke et al., 2008; Rustom et al., 2004; Sowinski et al., 2008). Leukocytes, their leukemic counterparts and bone marrow stromal cells have all been reported to form TNTs *in vitro* (Andresen et al., 2013; Chauveau et al., 2010; Matula et al., 2016; Omsland et al., 2017; Onfelt et al., 2004; Polak et al., 2015; Reichert et al., 2016). TNTs might represent a mechanism for chemo resistance in e.g. by transport of oncoproteins as shown between T and B cells, by transfer of mitochondria from endothelial cells to chemotherapy exposed cancer cells, or by induced drug-efflux in aggressive forms of pancreatic carcinoma (Ahmad et al., 2014; Desir et al., 2018; Pasquier et al., 2013; Rainy et al., 2013; Wang and Gerdes, 2015). The impact of TNTs *in vivo* is so far not well characterized, but it has been described to connect myeloid cells in the cornea of mouse (Chinnery et al., 2008; Seyed-Razavi et al., 2013) and in resected solid tumors from patients with malignant pleural mesothelioma and lung adenocarcinoma *in vivo* (Lou et al., 2012).

Here, the role of BCR-ABL1 on TNT formation in CML cells has been characterized. We found low TNT numbers in CML cells, while treatment with IFNα or the ABL1 inhibitor nilotinib swiftly induced TNT formation involving β1-integrin.

## Results

### TNT formation in Kcl-22 cells is increased following treatment with IFNα or the tyrosine kinase inhibitor nilotinib

In order to investigate the presence of TNTs between CML cells, primary bone marrow CML cells were cultured for 24 h on fibronectin coated surfaces and TNTs were quantified as earlier described for acute myeloid leukemia (AML) cells (Omsland et al., 2017). When we compared number of TNTs/100 cells in bone marrow cells derived from four different patients diagnosed with CML (P1-P4), very few TNTs were detected and the cells appeared mobile and morphologically spherical (Fig. 1A and Fig S1).

**Fig 1:**
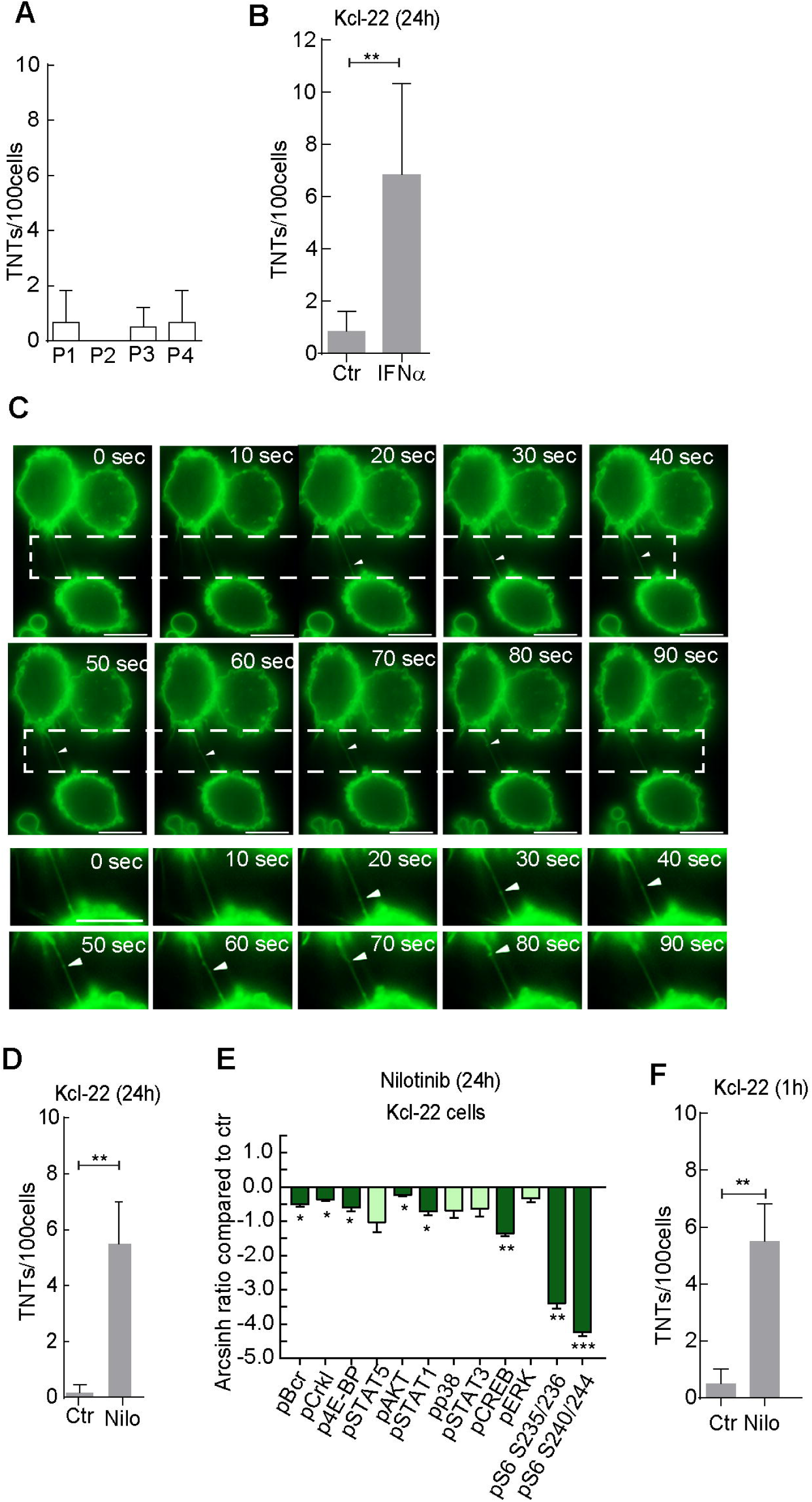
CML therapy influence TNT formation in CML cells. (A) TNT quantification of bone marrow samples from 4 different CML patients, results are presented as number of TNTs/100 cells from the average of duplicates. (B) Number of TNTs were quantified in Kcl-22 (memGFP) cells treated with 100 U/ml of IFNα for 24 h compared to untreated (Ctr) (C) Time-lapse of Kcl-22 cells (memGFP) treated for 1 h with IFNα where images were captured every 10^th^ second for a total of 120 seconds. Arrow heads show movement of memGFP along the TNT structure over time. (D) Kcl-22 (memGFP) cells were untreated (Ctr) or treated with 100 nM nilotinib (Nilo) for 24 h. (E) Mass cytometry analysis of down-stream signaling pathways of BCR-ABL1 in Kcl-22 cells treated with nilotinib (100 nM) for 24h. Results are illustrated by fold changes relative to control (all gated for live cells) based on calculated Arcsinh Ratio of Medians, median from three independent experiments are shown. (F) Kcl-22 cells were untreated (Ctr) or treated with 100 nM nilotinib (Nilo) for 1 h. Scale bar = 10 μm. For all displayed graphs: Mean ±standard deviation (s.d.) used together with unpaired t-tests (P**<0.005, n.s= not significant). All TNT quantifications were performed at least three independent times unless otherwise noted. Fluorescence microscopy was performed by the use of AxioObserver Z1 fluorescence microscope (Carl Zeiss, Inc, Thornwood, NY) with Alpha Plan Apochromat 63X/1.4 NA Oil DICIII.

To further investigate the effect of drug treatment on TNT formation in CML cells, the CML cell line Kcl-22 was examined before and after treatment with standard CML therapeutics. To enable live imaging of TNTs the cell line were stably transduced to express a cellular membrane localized GFP (memGFP). Similar to the primary patient cells, the Kcl-22 cells also demonstrated very low numbers of TNTs of 0.8 TNTs/100 cells, however, following 24 h IFNα treatment (100 U/ml) resulted in an increase of TNTs to 6.8 TNTs/100 cells (Fig. 1B). Time-lapse microscopy following 1 h treatment with IFNα (100 U/ml) demonstrated GFP positive dots moving along the TNTs, from one cell to another, indicating function as transport devices (Fig. 1C and Movie S1). Next, we treated Kcl-22 for 24 h with pre-apoptotic concentration of the Abl1 tyrosine kinase inhibitor (TKI) nilotinib (100 nM) and quantified the number of TNTs compared to untreated cells. This also resulted in induced TNT formation (Fig 1D). Cell viability after nilotinib treatment was investigated by Hoechst staining, where 24 h treatment induced 10% cell death. Inhibition of BCR-ABL1 signaling by nilotinib (100 nM) for 24 h was verified by single cell mass cytometry analysis of phospho-specific antibodies (Gullaksen et al., 2017). The nilotinib treatment resulted in a reduction in phosphorylation of CRKL, STAT5 and CREB among others, in Kcl-22 cells (Fig. 1E and Fig. S2B). The TNT inducing effect of nilotinib was apparent already after one hour treatment when we treated the Kcl-22 cells for one hour with nilotinib (Fig. 1F).

### Nilotinib treatment induces TNTs in an actin dependent manner

Immunofluorescence microscopy of nilotinib-treated Kcl-22 cells (1 h) revealed the presence of F-actin in the TNTs and the absence of β-tubulin (Fig. 2A). A morphological change was observed for the nilotinib-treated Kcl-22 cells, from spherical semi-attached cells to more spread-out and firmly attached cells (Fig. 2A). The critical role of F-actin in these TNTs was further examined by treating the cells with the actin polymerization inhibitor cytochalasin D (CytD (Casella et al., 1981)). Kcl-22 cells were treated for 24 h with nilotinib (100 nM) and quantified for TNTs before treatment with CytD (2 μM) for 20 min followed by a second TNT quantification (Fig. 2B). This showed that the CytD treatment resulted in TNT collapse and less prominent cell stretching (Fig. 2B, C). These data demonstrate that inhibition of BCR-ABL1 by nilotinib induces the formation of TNTs in actin polymerization dependent manner.

**Fig 2:**
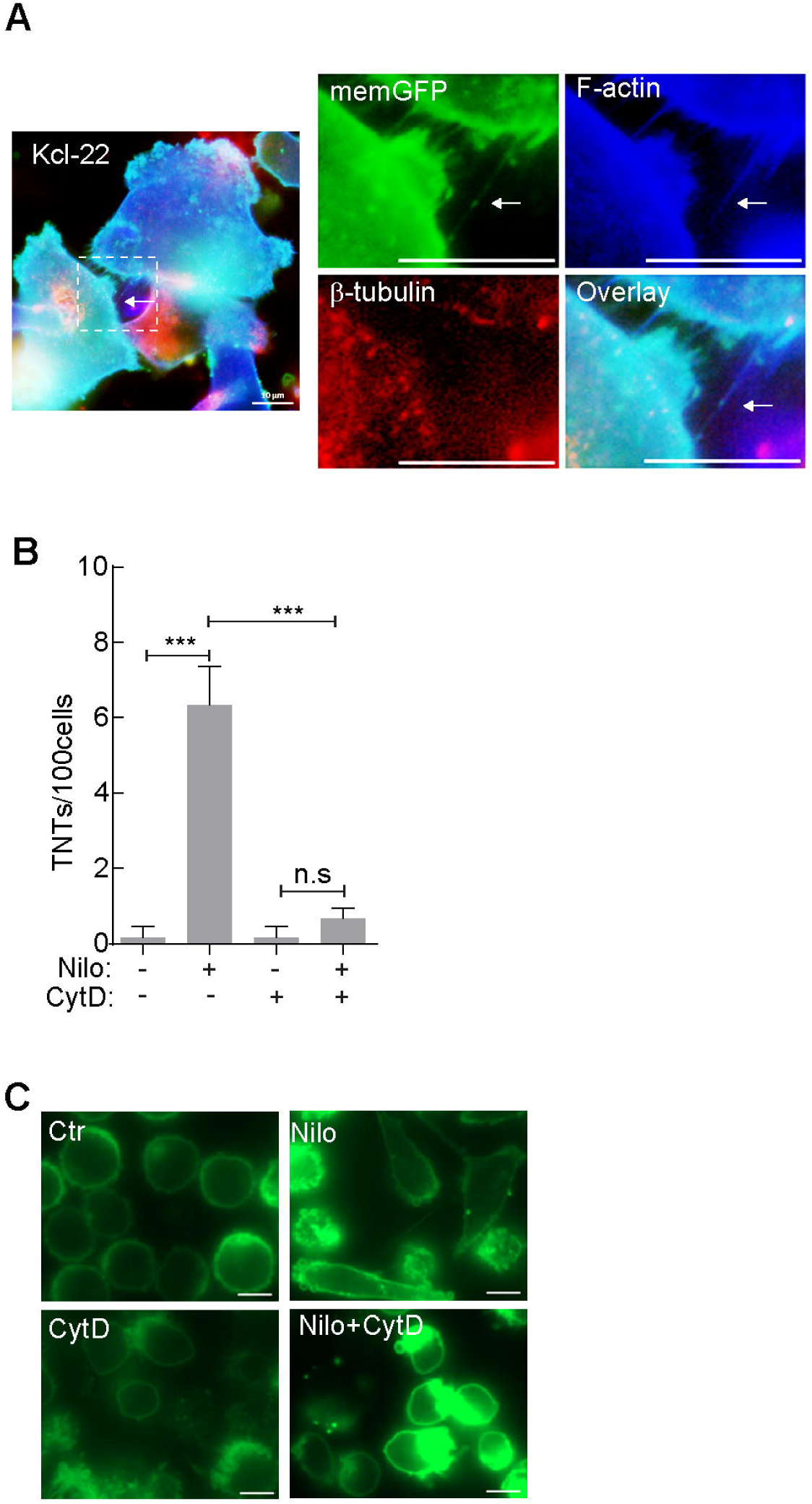
Nilotinib induces TNTs in an actin dependent manner. (A) Kcl-22 cells treated with 100 nM nilotinib 1 h, fixed with 4% PFA and stained with phalloidin AF350 followed by anti-β-tubulin staining. Representative images of three independent experiments are shown. Scale bars = 10 μm. (B) Kcl-22 cells were untreated (Ctr) or treated with 100 nM nilotinib (Nilo) for 24 h and TNT quantification was performed before and after addition of CytochalasinD (CytD, 2 μM) for 20 min, at 37°C. (C) Representative fluorescence images from three independent experiments performed in duplicate of Kcl-22 (memGFP) cells with no treatment (Ctr) or treatment with nilotinib (Nilo), cytochalasin D (CytD) or nilotinib (Nilo) followed by cytochalasin D (CytD). Scale bar = 10 μm. For all displayed graphs: Mean ±standard deviation (s.d.) used together with unpaired t-tests (P**<0.005, P***<0.001, n.s= not significant). All TNT quantifications were performed at least three independent times. Fluorescence microscopy was performed by the use of AxioObserver Z1 fluorescence microscope (Carl Zeiss, Inc, Thornwood, NY) with Alpha Plan Apochromat 63X/1.4 NA Oil DICIII.

### Expression of BCR-ABL1 results in reduced TNT formation and spherical cell shape

To further study the involvement of BCR-ABL1 in TNT formation, a doxycycline inducible BCR-ABL1 protein (Klucher et al, 1998) (p210) was introduced in Ba/F3 cells. Ba/F3 cells represent a well explored system for characterization of the oncogene function of BCR-ABL1, where expression of BCR-ABL1 allows Ba/F3 cells to proliferate independent of IL-3 (Daley and Baltimore, 1988). The induction of BCR-ABL1 expression by doxycycline was verified by immunoblotting and IL-3-independent proliferation (Fig. 3A, Fig. S2A). In the Ba/F3 cells, BCR-ABL1 expression resulted in a morphological change from mostly non-spherical and semi-attached cells to spherical and less firmly attached to the fibronectin coated plastic culture well (Fig. 3B). Interestingly, expression of BCR-ABL1 was also accompanied by down-regulation of TNTs (Fig. 3C). This down-regulation of TNTs was not due to the doxocycline treatment, since treatment of Ba/F3 cells transfected with an empty vector resulted in an increase in TNT formation rather than a decrease (Fig. 3C). Treatment of Kcl-22 cells with nilotinib (100 nM) or IFNα for 1 h resulted in the opposite change compared to the Ba/F3 cells from round non-attached cells to stretched firmly attached cells (Fig. 2A and 3D), suggesting that BCR-ABL1expression result in a spherical cell morphology, reduced attachment to fibronectin and TNT formation.

**Fig 3:**
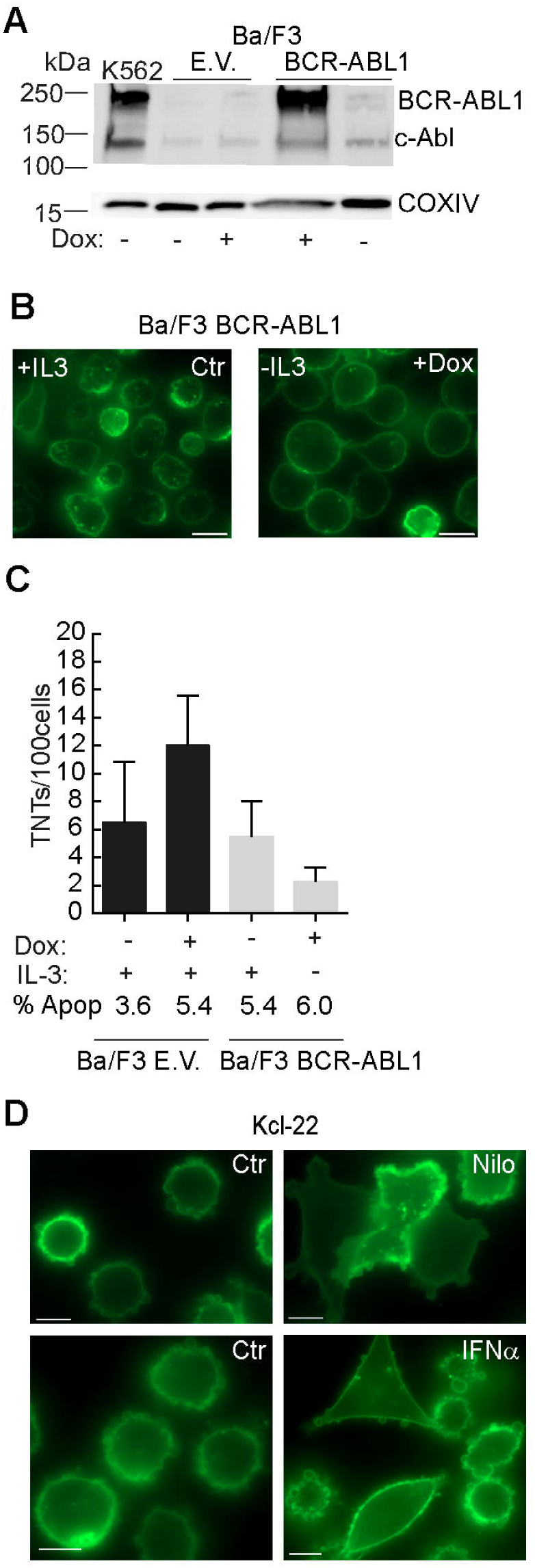
BCR-ABL1 effects of TNT formation and cell morphology. (A) Immunoblotting of Ba/F3 BCR-ABL1 doxycycline (Dox) inducible cells. Ba/F3 cells transduced with empty vector (E.V.) and Ba/F3 transduced with BCR-ABL1 were untreated or treated with 0.1 μM doxycycline for 24 h. Anti-cAbl antibody was used to verify BCR-ABL1 expression. K562 cells were used as positive control and COXIV as loading control. (B) Fluorescence microscopy of Ba/F3 BCR-ABL1 doxycycline inducible cells cultured in the presence or absence of IL-3 and with (+Dox) or without (Ctr) 0.1 g/ml doxycycline, cells were stained with WGA488. Scale bars: = 10 μm. (C) TNT quantification of Ba/F3 transduced with empty vector (black bars) and BCR-ABL1 doxycycline inducible Ba/F3 cells (grey bars) cultured in the presence (+) or absence (-) of IL-3 from 10% WEHI conditioned medium and with (+) or without 0.1 g/ml doxycycline (-), 1 μg/ml puromycin was present in the culture media in all conditions. (D) Fluorescence microscopy of Kcl-22 (memGFP) cells untreated (Ctr) or treated for 1 h with nilotinib (Nilo) or IFNα (100 U/ml). Scale bar= 10 μm. (E) Microscopy was performed using AxioObserver Z1 fluorescence microscope (Carl Zeiss, Inc, Thornwood, NY) with Alpha Plan Apochromat 63X/1.4 NA Oil DICIII. All data are presented as mean ±standard deviation (s.d.) and investigated for significance by unpaired t-tests: (P**<0.005). All experiments were performed three times except TNT quantification of Ba/F3 treated with doxocycline and incubated without IL-3 (n=2).

### TNT formation and increased cell surface adhesion induced by drug treatment

Cell adherence to fibronectin has been found to correlate with TNT formation and treatment of CML cells with IFNα and TKIs have been showed to increase cell adherence to fibronectin, described through a restoration of the β1 integrin by IFNα (Bhatia and Verfaillie, 1998; Dowding et al., 1991; Obr et al., 2014; Reichert et al., 2016). To study the role of cell adherence through β1 integrin in TNT formation, we pre-incubated Kcl-22 cells for 30 min with a blocking antibody against β1 integrin before 1 h treatment with either IFNα (100 U/ml) or nilotinib (100 nM). The control Kcl-22 cells, not pre-incubated with the β1 integrin blocking antibody, changed cell morphology and significantly changed the cell surface area (μm^2^) on the fibronectin coated culture wells following nilotinib treatment, whereas IFNα treatment only resulted in altered morphology without significant changes in cell surface area (Fig. 4A-B). Strikingly, these nilotinib and IFNα-induced changes in cell morphology were completely blocked by pre-incubation with the β1 integrin blocking antibody (Fig. 4C-D). When cell motility was measured by time-lapse microscopy, the IFNα and nilotinib-induced change in cell morphology was associated with a significant decrease in cell motility. Conversely, pre-treatment with the β1 blocking antibody resulted in increased cell motility (Fig. 4C-D) suggesting a direct connection between increased functionality of integrin β1 and cell adherence, here induced by IFNα and nilotinib leading to increased TNT formation.

**Fig 4:**
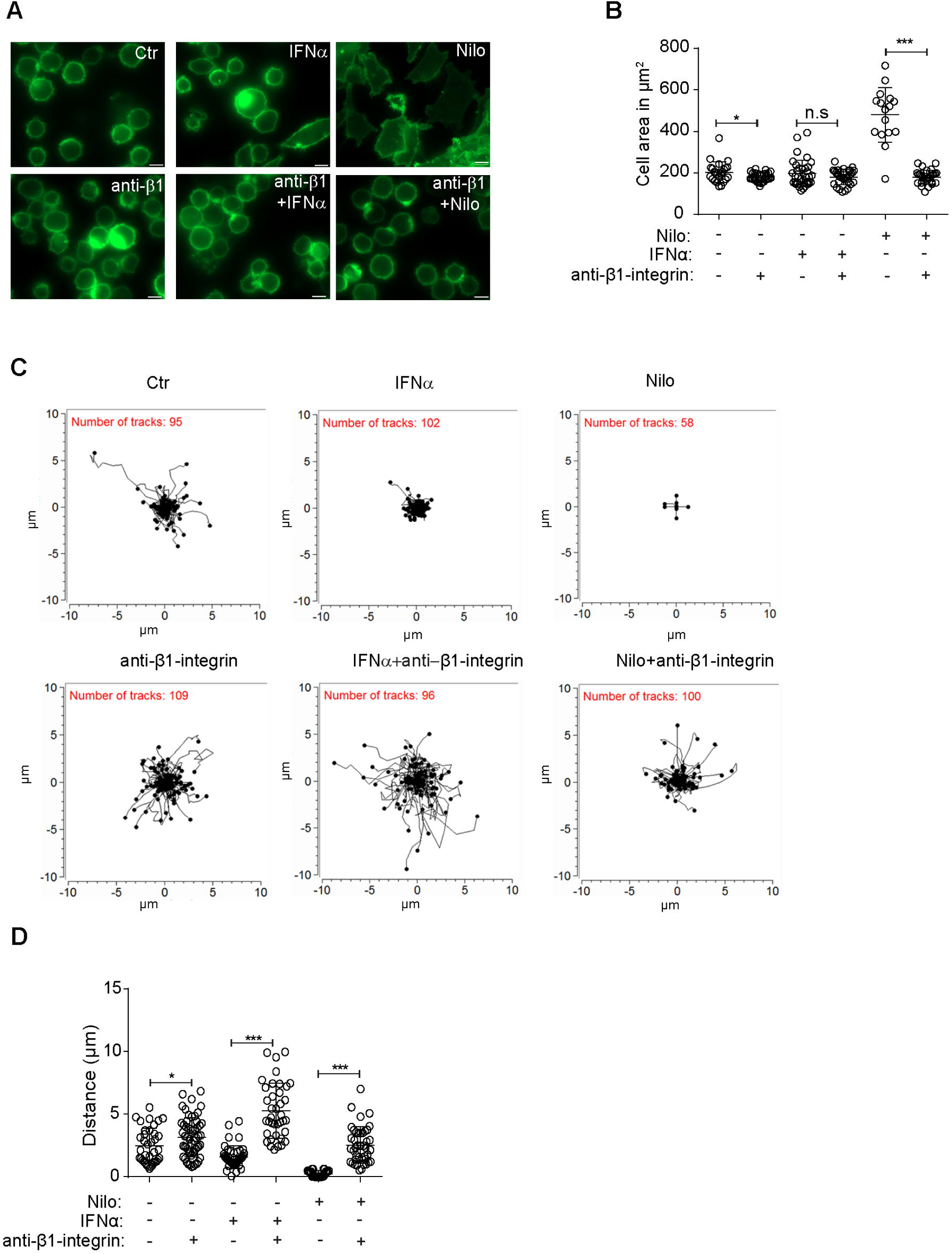
CML cell adherence to fibronectin enhances TNT formation and reduce cell mobility. (A)Kcl-22 (memGFP) cells were pre-treated for 30 min with anti-β1 integrin blocking antibody (10 μg/ml) before seeded in fibronectin-coated IBIDI wells and allowed to adhere for 3 h before treated for 1 h with nilotinib (Nilo) (1 μM) or IFNα (100 U/ml). Cells were investigated by fluorescence microscopy. Scale bars = 10 μm. (B) Cell area of experiments in (A) was measured manually using ImageJ. (C) Cells seeded on fibronectin were tracked for motility by live cell imaging and analyzed using using metamorph and measurements were calculated using Chemotaxis and Migration (IBIDI) plugin in ImageJ. (D) Statistical analysis of motility of the Kcl-22 cells following the different treatment conditions. Significant changes were calculated using unpaired Mann-Whitney test. Mean ±standard deviation (s.d.) (P*<0.05, P***<0.001, n.s= not significant). Results are presented as mean ±standard deviation (s.d.) and significance investigated by the use of unpaired t-tests (P***<0.001, n.s= not significant).

## Discussion

TNT is a dynamic 50-200 nm structure consisting of plasma membrane and F-actin, but with elusive understanding of how its formation is regulated (Zaccard et al., 2016). Since the BCR-ABL1 fusion protein in CML has a strong impact on F-actin and simultaneously affects various signaling pathways (Van Etten et al., 1994) we examined the effect of BCR-ABL1 on TNT formation. Both bone marrow derived BCR-ABL1 positive cells from CML patients and the CML cell line Kcl-22 displayed low numbers of TNTs compared to acute myeloid leukemia cells and other cancer cells (Hase et al., 2009; Omsland et al., 2017; Reichert et al., 2016). One possible explanation for the low TNT numbers could relate to the observation that CML cells adhere poorly to the bone marrow stroma (Gordon et al., 1987), consequently resulting in interrupted cellular TNT communication. TNT formation between cells *in vitro* is highly dependent on adherence, and culturing leukocytes on a supportive layer of mesenchymal stem cells (MSCs) or fibronectin increase TNT formation (Osteikoetxea-Molnar et al., 2016; Reichert et al., 2016). IFNα was the first effective CML therapy and is now being re-evaluated in combination with TKIs like dasatinib (Apperley, 2015; Hjorth-Hansen et al., 2016). Interestingly, one of the proposed mechanisms for the efficacy of IFNα in treatment of CML patients was through its ability to restore adhesion of CML cells to the bone marrow stroma (Dowding et al., 1993). Similarly, TKI treatments result in increased CML cell adherence to fibronectin (Obr et al., 2014). Together, these observations suggest that restoration of adherence in CML cells could be central to successful treatment of CML patients.

Interestingly, both increased adhesion and change in morphology was observed in the CML cell line Kcl-22 after treatment with nilotinib or IFNα accompanied by a significant increase in TNT formation (Fig. 1B, D and Fig. 3D). Evidence for an involvement of BCR-ABL1 in TNT formation was obtained using the doxycycline-inducible system of BCR-ABL1 expression in the Ba/F3 cells. BCR-ABL1 induction caused these cells to appear more morphologically spherical compared to the Ba/F3 control cells (Fig. 3B). This confirmed observations by others where BCR-ABL1 expression in Ba/F3 cells induced cell detachment, and increased motility (Salgia et al., 1997).

To verify the importance of cell adhesion in TNT induction we incubated Kcl-22 cells with an integrin β1 blocking antibody before treatment with IFNα or nilotinib. Indeed, we found that the cell adhesion effect by the two therapeutics were dependent of β1 integrin (Fig. 4C). The Kcl-22 cells showed increased mobility and a more spherical morphology after pre-incubation with the integrin β1 blocking antibody (Fig. 4D). Together with the results obtained in the Ba/F3 cell model system, this supports a hypothesis where these CML cells form few TNT structures when adhering poorly to fibronectin as a consequence of showing a spherical appearance.

Taken together, we find that TNTs were induced in Kcl-22 cells following IFNα and nilotinib treatment as a result of increased cell adhesion after restoration of the β1-integrin function. TNT communication might be an important factor of the successful treatment of CML which merits further investigation in the future.

## Materials and Methods

### Cell lines

Kcl-22 and Ba/F3 cells (ATCC and DSMZ, mycoplasma tested while experiments were carried out) were cultured according to provider’s instructions. RPMI-1640 medium was supplemented with 10% FBS, 1% L-glutamine (2mM) and 1% (1.0 U/ml) penicillin and streptomycin (5mM) (Sigma-Aldrich). The RPMI-1640 medium for the IL-3 dependent Ba/F3 cells were additionally supplemented with 10% conditioned medium from WEHI3B cells (mouse myelomonocytic cell line) known to secrete high amounts of IL-3 (Lee et al., 1982). The WEHI3B cells were grown to confluency in a T75 flask with complete IMDM medium (containing 10% FBS, 1% Pen-Strep and L-glutamine), and cultured for 2-3 days before the supernatant was centrifuged at 1500 RPM for 10 min and sterile filtered through a 0.2 μm filter.

### Mem-GFP transduced cells

The memGFP-Kcl-22 cells were generated by transducing the cells with ready-to-use lentiviral particles expressing a membrane localization signal (20 amino acids of the N-terminal part of neuromodulin, containing a palmitolylation signal) fused to GFP; rLV-EF1-AcGFP-Mem-9 (Takara, rV2.1A1.1941 C2) according to the provider’s instructions. The transduced cells were sorted using BD FACS Aria SORP at the Flow Cytometry Core Facility, Department of Clinical Science, University of Bergen, Norway.

### Primary cells

The study was conducted in accordance with the Declaration of Helsinki and approved by the local Ethics Committee (Regional Ethics Committee West projects 2012/2245 and 2012/2247, University of Bergen, Norway). Blood and bone marrow samples from consecutively diagnosed CML patients were collected after informed consent and were processed by density gradient separation (Lymphoprep, Axis-Shield, Oslo, Norway) (Bruserud et al., 2001).

### Doxycycline inducible Ba/F3 cells

BCR-ABL1 (P210) was cloned into pcDNA3 (Adgene) after EcoRI digestion. The orientation and sequence was verified by PCR. This was further sub-cloned into the EcoRI site of PLVX-tetOne-Puro (from the Lenti-X Tet-One Inducible Expression Systems). Wild type Ba/F3 (kind gift to Prof. Enserink from Prof. Gordon Mills laboratory, Houston, Texas, USA) was transfected with 2 μg of the PLVX_tetOne_BCR-ABL1 plasmid or PLVX_tetOne_empty vector by electroporation (Amaxa biosystems nucleofector II: program U20) using Ingenio Electroporation solution (catalog number MIR 50114). Transfected cells were cultured in medium for 24-72 h before selection with 1 μg/ml puromycin. Puromycin resistant clones were sorted and grown independently; cells were continually cultured in medium with puromycin to maintain selection pressure. 0.1 μg/ml doxycycline was added to induce expression of BCR-ABL1.

### Antibodies and reagents

The following primary antibodies were used for immunofluorescence and/or immunoblotting: anti-β-tubulin (clone TUB 2.1, Sigma-Aldrich, 1:1000), anti-COX IV (ab16056, 1:2500), anti-cAbl ((24-11) sc-23, Santa Cruz Biotechnology, 1:1000), anti-integrin β1 blocking antibody [P5D2] (ab24693, Abcam). Secondary antibodies used for immunofluorescence or immunoblotting; Alexa Fluor^©^ 488- or 594-conjugated goat-anti-mouse (Invitrogen, 1:5000), horseradish peroxidase (HRP)-conjugated goat anti-rabbit/mouse (Jackson Immunoresearch, 1:10 000). The following were used for actin and membrane staining; AlexaFluor^©^ 350-conjugated phalloidin and wheat germ agglutinin (WGA) –Alexa Fluor^©^ 594 or 488 (Invitrogen) as previously described (Omsland et al., 2017). Tyrosine kinase inhibitor: Nilotinib (Selleckchem). Interferon alpha (IFNα) (Intron A from MSD), Cytochalasin D (Sigma-Aldrich), doxycycline (Doxyferm, Nordic Drugs AB, Limhamn), puromycin (Sigma-Aldrich), Bovine serum albumin (BSA) fraction V (Roche), fibronectin (Sigma-Aldrich). The determination of the concentrations of new antibodies was carried out in the laboratory when they arrived.

### TNT identification and quantification

A TNT in this study is defined as a thin straight structure, ≤ 200 nm in diameter, minimum 5 μm in length, hovering above the substratum, connecting two cells. TNTs were distinguished from cytoplasmic bridges, which appear following cell division, by the lack of a midbody clearly visible by differential interference contrast and/or staining of cellular membranes (Omsland et al., 2017). 8-well μ-slides (Ibidi GmbH) were pre-coated with fibronectin (10 μg/ml, F2006, Sigma-Aldrich) for 30 min at 37°C before washing with saline. 70000 cells were seeded per well and incubated overnight under physiological conditions. Primary CML cells were seeded in DMEM medium containing 20% FBS overnight and stained with wheat germ agglutinin conjugated with alexa fluor 488 or 594 (1.67 μg/ml) as previously described (Omsland et al., 2017). Cells were examined live by fluorescent light microscopy (Zeiss Axio Observer Z1 with AxioVision 4.8.2 or Zen software) using a 63X/1.4 NA Oil DICIII objective, heat block (37°C) and standard air conditions. 100 cells per well were counted following a fixed counting pattern with 5-6 cells examined per vision field. The result is described as number of TNTs/100 cells meaning the total number of TNTs (one TNT always connects two cells) among 100 cells counted. For further details see Supplementary Figure in Omsland et al (Omsland et al., 2017). Cell viability was monitored by Hoechst 3342 (Sigma) staining as previously described (McCormack et al., 2012).

### Blocking of integrin β-1

Cells were cultured in a 0.7×10^6^ cells/ml density in a 6-well plate. Cells were incubated in medium without or with 10 μg/ml of anti-integrin beta 1 [P5D2] antibody for 30 min before seeded to fibronectin pre-coated μ-slides (Ibidi GmbH). Cells were incubated for 3 h to allow attachment before treatment with 1 μM nilotinib (nilo) or 100 U/ml IFNα 1 h prior to examination by live microscopy.

Measuring of cell area was performed manually using ImageJ: Images were analyzed as 8-bit files using FFT Bandpass Filter, threshold was set manually and adjusted until cells were distinguished from the background>convert to mask>fill holes>cells in close proximity were then distinguished using watershed algorithm. Measuring of the cell area was performed using the measure tool under the region of interest manager tool and single cells were selected using the wand tool.

Tracking of cells was performed using metamorph and the chemotaxis and migration (Ibidi GmbH) plugin to ImageJ was performed to calculate accumulated distance and to make trajectory plots as described in (Hurley et al., 2013).

### Immunofluorescence

The F-actin and microtubule presence in TNTs was investigated in Kcl-22 cells (on 8-well μ-slides, Ibidi GmbH) fixed in 4% PFA in PBS and 0.2% glutaraldehyde in PBS for 20 min at room temperature (RT) followed by one wash with PBS, before permeabilized for 1 min using 0.2% Tween^©^ in PBS and washed twice with PBS. Cells were blocked with 0.5% Bovine Serum Albumin Fraction V (BSA) PBS for 20 min at RT and then incubated for 1 h at RT in the dark with 33nM AlexaFluor^©^ phalloidin, washed once with PBS and incubated with anti-β-tubulin antibody (1:200 in blocking solution) overnight at 4°C. Then cells were washed twice with PBS and incubated with Alexa-488 or 594 goat-anti-mouse antibodies (1:5000 in blocking solution) for 1h at RT, before washed twice with PBS and examined by fluorescence microscopy. Cells not expressing memGFP were stained with wheat germ agglutinin (WGA) conjugated with Alexa 488 or 594 for 8 min followed by one wash with PBS before examined by microscopy and manual quantification of TNTs.

### Immunoblotting

Cells were lysed and analyzed by immunoblotting according to standard protocol (Shieh et al., 1999; Silden et al., 2013). Briefly, immunobloting was performed using precast gels from BioRad, transferred to PVDF membranes using Pierce G2 fast blotter (Thermo Scientific). Membranes were blocked for 1h at RT in 5% fat-free drymilk or 5% BSA in TBST, incubated with primary antibody at 4°C overnight. Membranes were washed with TBST followed by incubation for 1 h with secondary antibody ((HRP)-conjugated goat-anti-rabbit/mouse) was diluted 1:1000 in 5% drymilk in TBST and washed with TBST before developed using SuperSignal West pico or femto (Thermo Fisher Scientific). Developed immunoblots were detected and captured by ImageQuant LAS 4000 (GE Healthcare Life Sciences).

### Mass Cytometry Barcoding

To reduce experiment variability, workload and antibody consumption, we used the commercially available metal barcoding kit from Fluidigm. Briefly, the cells from each sample were stained with a unique three-palladium isotope combination; three chosen from six available; Pd 102, Pd 104, Pd 105, Pd 106, Pd 108, Pd 110 (20 unique combinations available). After cell barcoding and washing according to the manufacturers’ recommendations, uniquely barcoded samples were pooled for further processing for mass cytometry analysis.

### Antibody staining

A pool of barcoded cells was stained with a panel of cell surface markers (30 minutes, RT) and permeabilized with methanol (−20^°^C). Further staining with intracellular phospho-specific antibodies (30 minutes, RT) followed. Cells were then washed and re-suspended in the buffer containing Iridium-intercalator (natural abundance iridium as pentamethylcyclopentadienyl-Iridium (III)-dipyridophenazine), which intercalates into the DNA (1 hour, 4^°^C), before washed and pelleted by centrifugation. Immediately prior to data acquisition cells were re-suspended to a final concentration of approximately 5 × 10^5^ cells/mL in MaxPar water (Fluidigm) containing normalization beads (1:10 dilution, Fluidigm) and analyzed on a Helios mass cytometer (Fluidigm), placed in the Flow Cytometry Core Facility of Bergen, University of Bergen.

### Single cell discrimination and barcoding de-convolution

Using the normalization beads and the normalization software, any drift in the data resulting from loss of detector sensitivity was abrogated. An automatic barcode deconvolution algorithm developed by Zunder *et al* 2015 (Zunder et al., 2015) was used to identify each uniquely barcoded sample. Further discrimination and gating of single cells was achieved by plotting all events by DNA-content (Ir 191 or Ir 103) versus Event Length (number of pushes). Together, barcode deconvolution and gating of cells on DNA content versus event length, is an effective filter for removal of doublets and identification of single cells. Finally, cleaved Caspase 3 readily discriminated between apoptotic and non-apoptotic cells, where non-apoptotic cells were used for statistical analysis.

**Table 1.** Antibody panel for mass cytometry analysis

| A.m.u | Metal | Epitope | Clone | Vendor |
| --- | --- | --- | --- | --- |
| 102 | Pd | Metal Barcode Channel #1 | N.A. | Fluidigm |
| 104 | Pd | Metal Barcode Channel #2 | N.A. | Fluidigm |
| 105 | Pd | Metal Barcode Channel #3 | N.A. | Fluidigm |
| 106 | Pd | Metal Barcode Channel #4 | N.A. | Fluidigm |
| 108 | Pd | Metal Barcode Channel #5 | N.A. | Fluidigm |
| 110 | Pd | Metal Barcode Channel #6 | N.A. | Fluidigm |
| 89 | Y | CD45 | HI30 | Fluidigm |
| 141 | Pr | pBCR Y177 | Polyclonal | Cell Signaling Technologies |
| 142 | Nd | Caspase 3 Cleaved | D3E9 | Fluidigm |
| 143 | Nd | pCrkL [Y207] | Polyclonal | Fluidigm |
| 149 | Sm | p4E-BP1 | 236B4 | Fluidigm |
| 150 | Nd | pStat5 [Y694] | 47 | Fluidigm |
| 153 | Eu | pStat1 [Y701] | 58D6 | Fluidigm |
| 154 | Sm | pAbl Y245 | 73E5 | Cell Signaling Technologies |
| 156 | Gd | p-p38 [T180/Y182] | D3F9 | Fluidigm |
| 158 | Gd | pStat3 [Y705] | 4/P-STAT3 | Fluidigm |
| 165 | Ho | pCREB [S133] | 4/P-STAT3 | Fluidigm |
| 167 | Yb | pERK 1/2 [T202/Y204] | D1314.4E | Fluidigm |
| 172 | Yb | pS6 [S235/S236] | N7-548 | Fluidigm |
| 176 | Yb | pS6 [S240/S244] | D68F8 | Cell Signaling Technologies |
| 191 | Ir | DNA | N.A. | Fluidigm |
| 193B | Ir | DNA | N.A. | Fluidigm |

### Statistical analysis

Differences between two groups were analyzed by two-tailed unpaired T-test using GraphPad Prism 6 Version 6.03. F-test was performed to verify that the internal variance in the groups were not significant. Significant difference was considered by a P-value <0.05. For cell area and cell movement unpaired Mann Whitney tests were performed.

## Acknowledgements

We thank Dr. André Sulen for help with sorting the Kcl-22-mem-GFP cells and Calum Leitch and Dr. Genoveffa Franchini for helpful feedback on the manuscript. We thank Dr. Tatiana Karpova for assistance with the analysis of cell movement.

